# A dynamic microbial community with high functional redundancy inhabits the cold, oxic subseafloor aquifer

**DOI:** 10.1101/171934

**Authors:** Benjamin J. Tully, C. Geoff Wheat, Brain T. Glazer, Julie A. Huber

## Abstract

The rock-hosted subseafloor crustal aquifer harbors a reservoir of microbial life that may influence global marine biogeochemical cycles. Here we utilized genomic reconstruction of crustal fluid samples from North Pond, located on the flanks of the Mid-Atlantic Ridge, a site with cold, oxic subseafloor fluid circulation within the upper basement. Twenty-one samples were collected during a two-year period at three different depths and two locations with the basaltic aquifer to examine potential microbial metabolism and community dynamics. We observed minor changes in the geochemical signatures over the two years, yet a dynamic microbial community was present in the crustal fluids that underwent large shifts in the dominant taxonomic groups. An analysis of 195 metagenome-assembled genomes (MAGs) were generated from the dataset and revealed a connection between litho- and autotrophic processes, linking carbon fixation to the oxidation of sulfide, sulfur, thiosulfate, hydrogen, and ferrous iron in a diverse group of microorganisms. Despite oxic conditions, analysis of the MAGs indicated that members of the microbial community were poised to exploit hypoxic or anoxic conditions through the use of microaerobic cytochromes and alternative electron acceptors. Temporal and spatial trends from the MAGs revealed a high degree of functional redundancy that did not correlate with the shifting microbial community membership, suggesting functional stability in mediating subseafloor biogeochemical cycles.

## Introduction

The largest actively flowing aquifer system on Earth is the fluids circulating through oceanic crust underlying the oceans and sediments (Sclater *et al.*, 1980; Carol A Stein and Seth Stein, 1994; Johnson and Pruis, 2003). The movement of water through the aquifer serves as a vital conduit for exchange of both microorganisms and nutrients between the ocean basins and the subseafloor and offers a route by which organisms can extract energy from the fluids and rocks beneath the seafloor (Orcutt *et al.*, 2013; Meyer *et al.*, 2016). Our understanding of life within the marine crustal aquifer has largely been shaped by studies of anaerobic and thermophilic organisms in warm ridge flank environments (Cowen *et al.*, 2003; Huber *et al.*, 2006; Jungbluth *et al.*, 2013; 2016) and crustal-source basalts exposed at the seafloor (Lysnes *et al.*, 2004; Mason *et al.*, 2009; Santelli *et al*., 2009; Lee *et al.*, 2015). However, much of the microbial interaction with the crustal aquifer occurs within the seafloor at sites where cold, oxygenated deep ocean waters circulate through basaltic crust, entering and exiting through seafloor exposures (Andrew T Fisher and Wheat, 2010; Edwards *et al.*, 2012). Therefore, despite advancing knowledge about microbial life in the subseafloor, our understanding is limited relative to which microorganisms live in the rocky oceanic crust, what hydrogeologic processes control subsurface fluid circulation, how these organisms harness energy in this environment, and the overall contribution to marine biogeochemical cycles is limited.

To more effectively study these prevalent ocean environments, several subseafloor observatories, termed Circulation Obviation Retrofit Kits (CORKs; Davis *et al.*, 1992; Wheat *et al.*, 2011), have been deployed in oceanic crust in part to allow for sampling and monitoring of the crustal aquifer (Wheat *et al.*, 2011). Two CORK observatories are installed at the well-studied site North Pond, an isolated sediment basin (8 km × 15km, ~4,484 meter water depth), just west of the Mid-Atlantic ridge on 7-8 million years old crust (22°45’ N, 46°05’ W; Edwards *et al.*, 2010). At North Pond, seawater circulates between the crust and the deep ocean through the exposed ridge flanks, while sediments within the basin act as an impermeable barrier that prevents seawater exchange.

Previous studies have sought to constrain the microbial community and its activity within the basaltic aquifer at North Pond. Measurements of carbon fixation activity on basalts recovered by ocean drilling (Orcutt *et al.*, 2015) were unable to detect quantifiable rates of activity at *in situ* temperatures (4°C), while additions of nitrate and ammonia to crustal rocks stimulated microbial growth (Zhang *et al.*, 2016). Modeling of the subsurface at North Pond suggests that hydrogen and ferrous iron likely have an important role in maintaining microbial biomass, with ferrous iron estimated to support ~10% of the microbial biomass (Bach, 2016). In support of this hypothesis, a *Marinobacter* isolate capable of iron oxidation was enriched from North Pond basalts (Zhang *et al.*, 2016). PCR-based assessments of the microbial community associated with the basalts from North Pond have shown that *Gammaproteobacteria* are the dominant phylogenetic group (Jørgensen and Zhao, 2016), while the presence of genes involved in the carbon fixation through the Calvin-Benson-Bassham cycle are more common than the reverse citric acid cycle (Orcutt *et al*., 2015).

Additional work examining the crustal fluids from the aquifer at North Pond has shown that the geochemistry of the fluids is nearly identical to the deep Atlantic bottom water (DABW), indicating a short residence time for seawater within the crustal aquifer at North Pond (Meyer *et al.*, 2016). However, basaltic formation fluids within the aquifer have concentrations of dissolved oxygen, silica, and dissolved organic carbon (DOC) that are different than those of the deep bottom water (Meyer *et al.*, 2016), and assessment of the crustal fluid microbial community through 16S rRNA gene and transcript sequencing, stable-isotope incubations, and metagenomics revealed that the aquifer community was active with a distinct community structure from bottom water. The community also had the capacity to perform both autotrophy and heterotrophy (Meyer *et al*., 2016), with low rates of activity detected using nanocalorimetry (Robador *et al.*, 2016).

Together, these initial studies show a diverse and distinct microbial community living in the oligotrophic, oxic, basaltic crustal aquifer at North Pond with relatively low levels of metabolic activity. However, little is known about the metabolic potential and community dynamics in this understudied environment. Here, we present genomic reconstruction of North Pond crustal fluid samples collected over a span of two years, providing 21 samples for a detailed examination of potential microbial metabolism and community interactions within this subseafloor aquifer. Our high-resolution analysis of hundreds of genomes reveals a temporally and spatially dynamic microbial community and provides new insights into microbially-mediated biogeochemical cycling within the crustal aquifer.

## Results

### Assessment of microorganisms in the subseafloor aquifer

The cold, oxic Mid-Atlantic subseafloor aquifer was sampled for geochemistry, cell quantification, and microbial DNA from two seafloor CORK installations at the North Pond site. Water samples were collected 10 times from hole U1382A over the course of two years, and Hole U1383C was sampled 9 times from three different depth horizons that had been sealed during These horizons are defined by packers that seal the borehole and limit vertical mixing, defining three distinct hydrologic zones based on formation properties (Edwards *et al.*; FIGURE 1; SUPPLEMENTARY FIGURE 1) Two additional background seawater samples were collected from Niskin bottles that were tripped approximately 50 meters off bottom, above the water-sediment interface (~4,450-m depth) in both 2012 and 2014. These samples proved a measure of bottom water properties using the same techniques employed on those from the crustal fluids (FIGURE 1).

**Figure 1.**
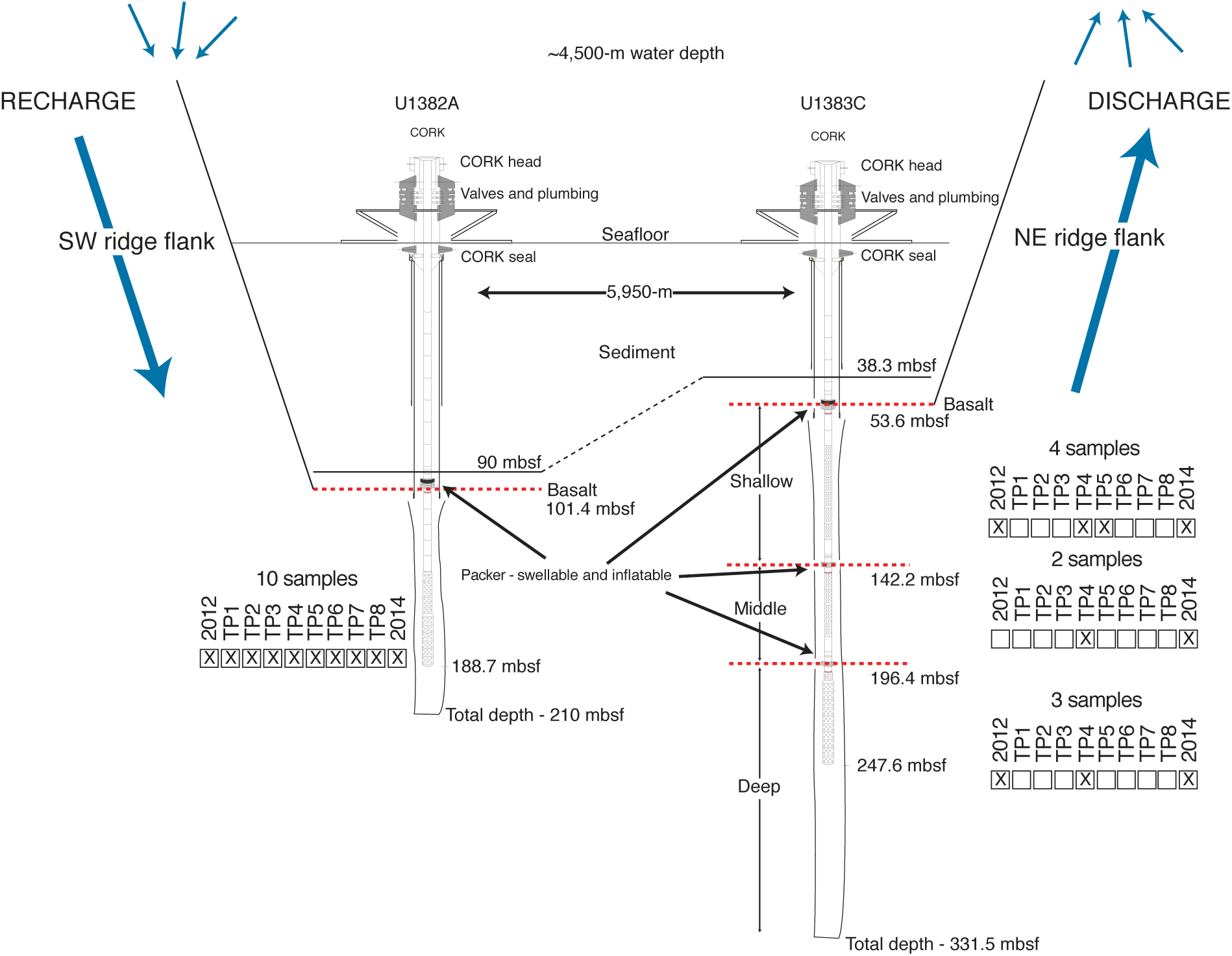
Idealized schematic of North Pond, CORK U1382A, and CORK U1383C. Hypothesized flow of entrained seawater through the crust is represented by blue arrows. Seafloor and sediment boundaries are represented by black line. Basalt boundaries are represent by red, dashed lines. For each CORK horizon, the number of metagenomic samples are indicated, including the relative time sampled. Abbreviations: CORK, Continuous Observation Retrofit Kits; TP, time point; mbsf, meters below seafloor. (Modified from Edwards *et al.*, 2010)

Cell counts in all 19 borehole samples ranged from 5 to 20 × 10^3^ cells mL^-1^ of crustal fluid, with no discernable change during the two year period (TABLE 1). Geochemical data from discrete samples collected in 2012 and 2014 indicated a minor increase in silica, whereas oxygen concentrations decreased slightly at all sampling horizons. Nitrate concentrations did not change.

**Table 1.**
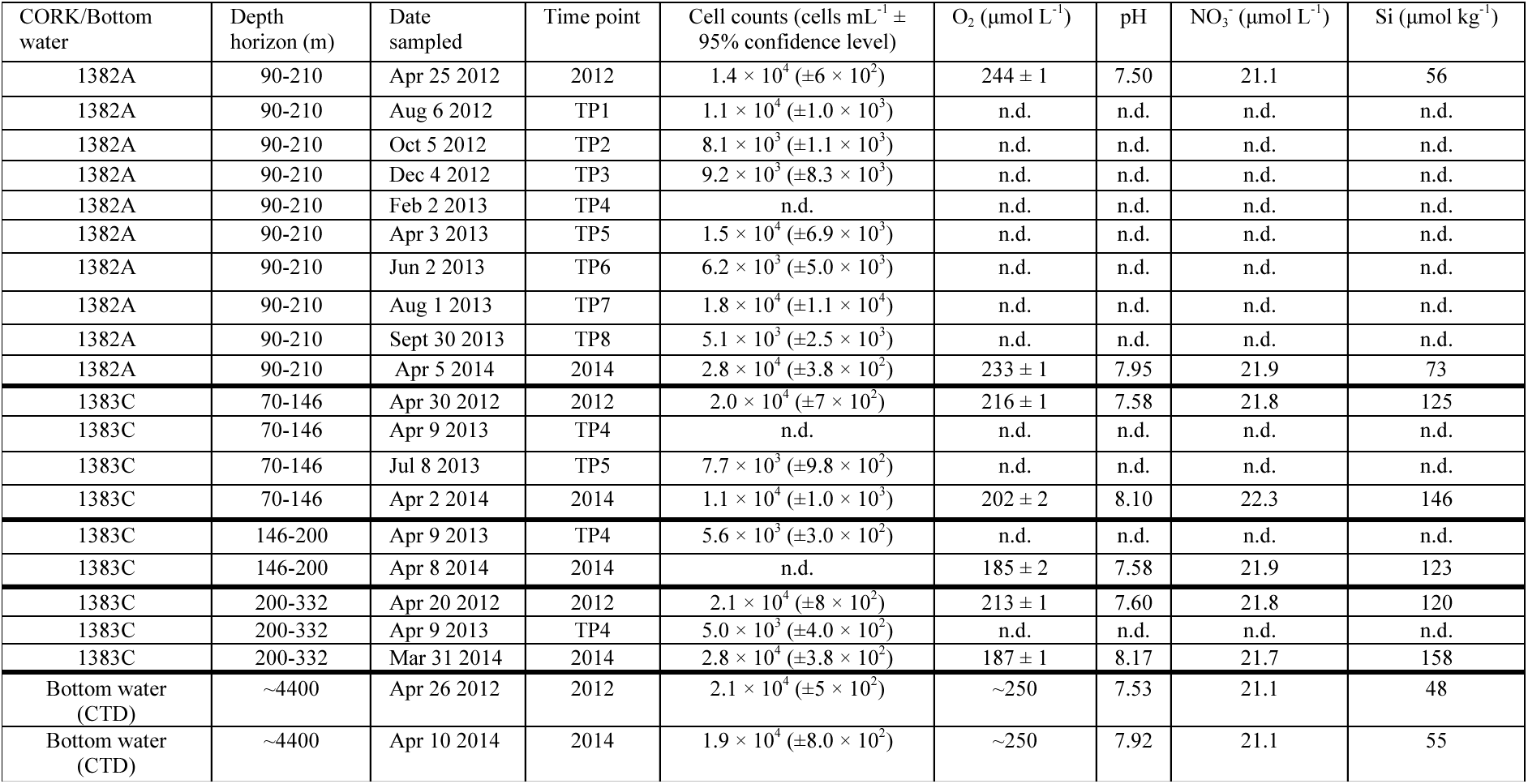
Collection details, cell enumeration, and inorganic chemical data for North Pond samples.

In total, 21 metagenomic samples were sequenced, generating 1.2 billion high-quality paired-end Illumina sequencing reads (SUPPLEMENARY DATA 1). 2,829 approximately full-length 16S rRNA gene sequences were reconstructed from the dataset (SUPPLEMENTARY DATA 2). The full-length 16S rRNA gene sequences in the metagenome (FIGURE 2) provide a snapshot of community composition in the samples, revealing a temporally and spatially dynamic community, with large shifts in the relative abundance of *Proteobacteria*, specifically in the *Epsilon*-, *Alpha*-, *Gamma*-, and *Deltaproteobacteria*, and the *Bacteroidetes.*

**Figure 2.**
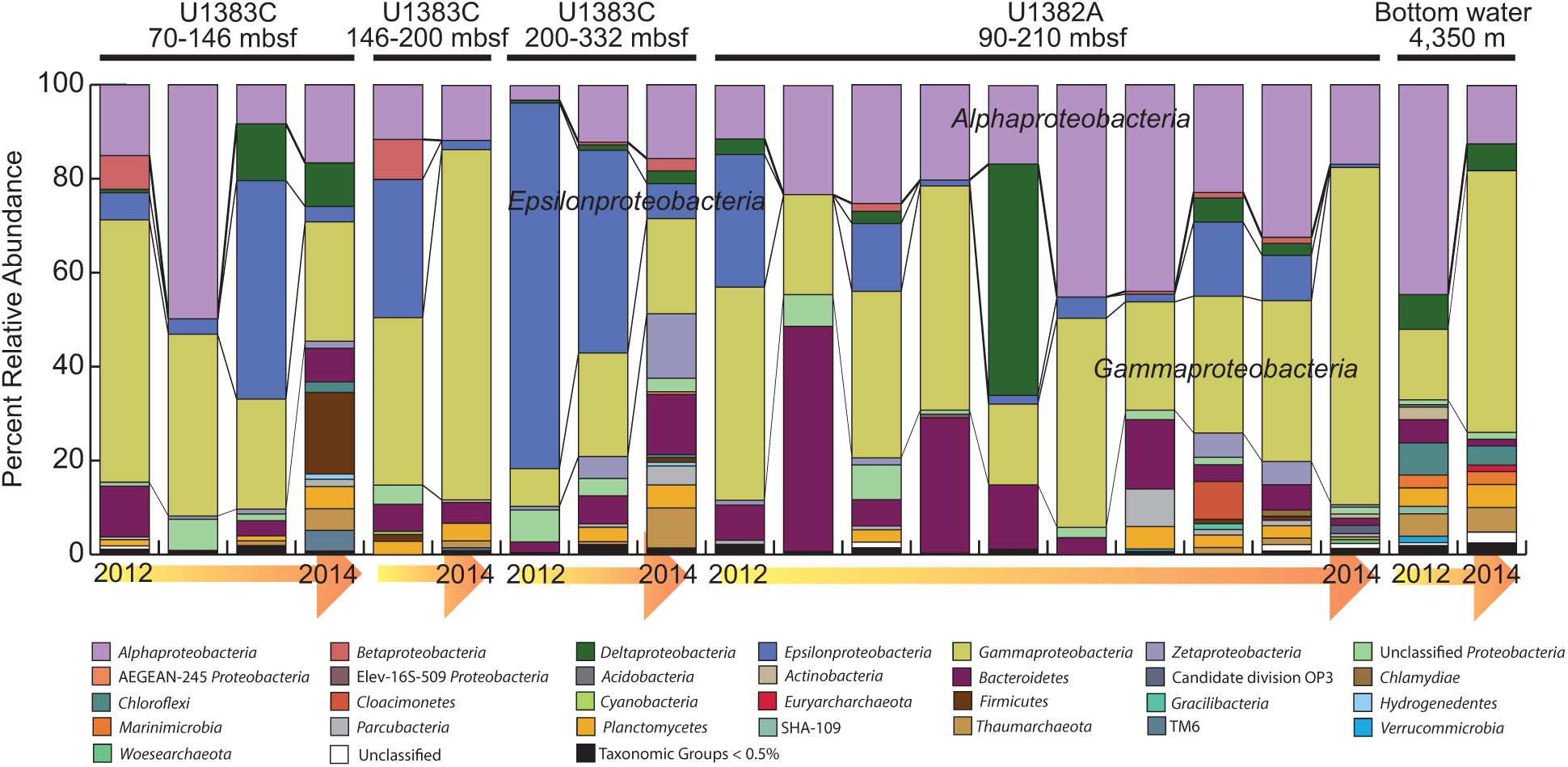
Microbial community structure in North Pond crustal fluids based on reconstructed full-length 16S rRNA gene sequences. Relative abundance for 16S rRNA gene sequences was combined at the Phyla (Class level for Proteobacteria) from holes U1383C and U13832A and the Deep Atlantic Bottom Water. Individual samples are grouped based on depth of sampling (shallow, middle, and deep in U1383C) and time (from initial sampling in 2012 to 2014 sampling time point).

After two rounds of assembly of the high-quality sequencing reads, 1.5 million contigs were produced. A subsection of contigs ≥5kbp in length (78,004 contigs; N50 = 25,932 bp; Total bp = 1.2 Gbp) were used to reconstruct microbial genomes (SUPPLEMENTARY DATA 3). 195 metagenome-assembled genomes (MAGs) were reconstructed and determined to be ≥50% complete (an additional 234 genome bins were identified that were 20-50% complete, though were not analyzed further). Throughout this manuscript, the term ‘genome’ will be used to refer to the 195 binned MAGs. The genomes were given the designation NORP, for **Nor**th **P**ond genome (NORP1-195). With the exception of two genomes (NORP4 and −5), all of the genomes had ≤10% cumulative contamination/redundancy (SUPPLEMENTARY DATA 3). The genomes recruited between 9-61% of the sequencing reads (mean = 37.3%) from the individual samples, with the lowest recruitment rate from the 2012 bottom water sample (SUPPLEMENTARY DATA 4).

140 MAGs had a sufficient number (≥8) of 16 ribosomal marker proteins to be included in a phylogenetic tree with genomes from IMG (Markowitz *et al.*, 2006) that represent the major bacterial Genera and/or Families (SUPPLEMENTARY DATA 5). The North Pond genomes were assigned to 20 Phyla, including all the lineages within the Proteobacteria (including *Acidithiobacillia*), the Candidate Phyla Radiation (CPR; Hug *et al.*, 2016), and the *Planctomycetes* (SUPPLEMENTARY FIGURE 2).

Based on the relative abundance of sequencing reads competitively recruited to each genome from each sample, the mean relative abundance for all genomes in all samples was 0.19% (median, 0.004%), and when examined closely, most genomes were ‘present’ in all samples at low abundance values (<0.05%; on average 151 of 195 genomes were below this threshold in each sample). NORP9 had the highest relative abundance (40.4%) in the 2012 U1383C deep sample (FIGURE 3; SUPPLEMENTARY DATA 9). Genomes were subjected to Bray-Curtis clustering based on the relative abundance values (SUPPLEMENTARY FIGURE 3). Several of the genomes (e.g., NORP125, −161, and −172) were cosmopolitan in the subseafloor crustal fluids, present in both holes, and at several time points and depths (FIGURE 3). Most of the genomes associated with the bottom water samples were not present in the crustal samples, although several genomes did have low abundances; specifically NORP160 and −164, both assigned to the *Nitrosopumilales*, and three additional genomes detected in the 2014 samples from U1382A and the middle section of U1383C. Generally, when genomes were grouped together, these groups were abundant in one or a few samples (e.g., NORP51, −54, and −55). These samples then tended to be linked either temporally or spatially (FIGURE 3). In several instances, a single organism becomes highly abundant, but is only present in a single sample (e.g., NORP6 or NORP73).

**Figure 3.**
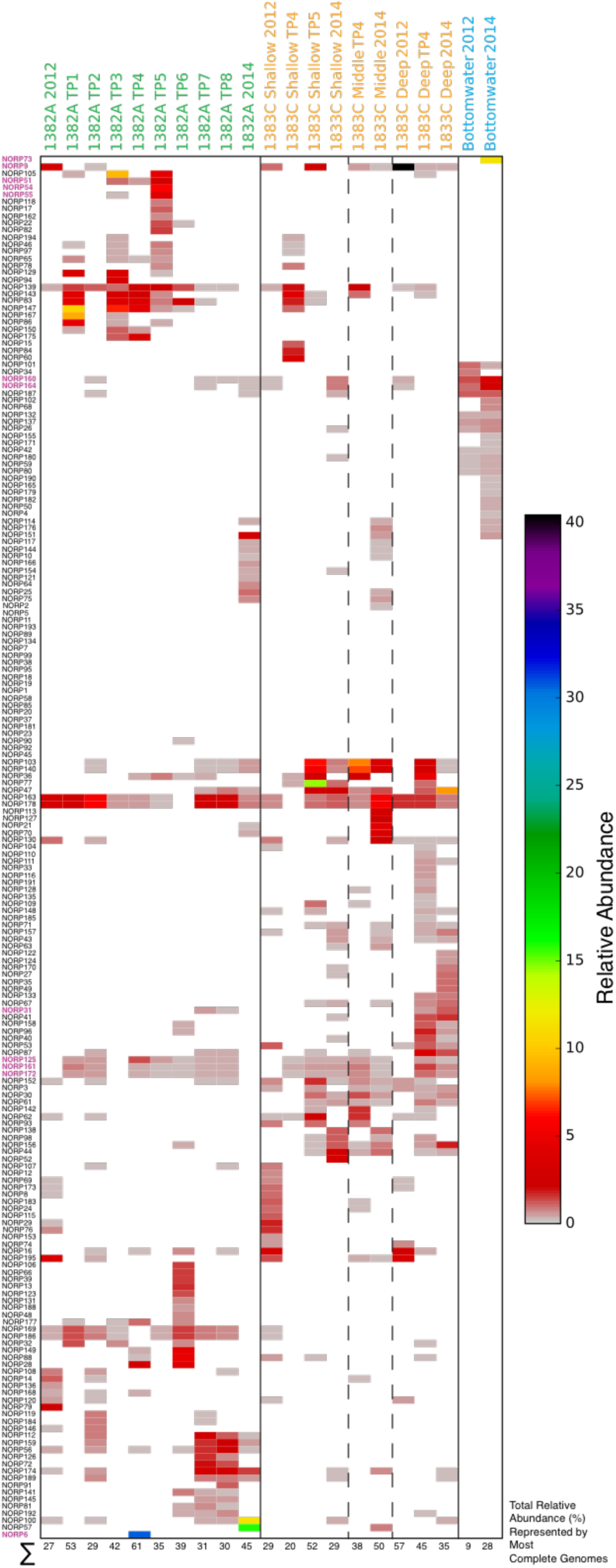
Heat map of relative abundances for the 195 high-quality genomes. Genomes are organized and clustered based on the Bray-Curtis similarity index (SUPPLEMENTARY FIGURE 3). Genomes explicitly mentioned in text are highlighted in bold and colored purple. Values at the bottom of the column represent the total observed relative abundance of genomes in that sample.

### Metabolic potential of metagenome-assembled genomes

From the genomes, 523,212 putative coding DNA sequences (CDSs) were identified, of which, 245,902 (47%) were annotated with a KEGG ontology (KO; Kanehisa, Sato, Kawashima, *et al.*, 2016) number/function (SUPPLEMENTARY FIGURE 4). Genomes were assessed for the presence of specific KO functions involved in numerous processes, including: carbon, nitrogen, sulfur, and hydrogen cycling, methanogenesis, motility, vitamin biosynthesis and transport, and fermentation pathways. Two of the 6 establish carbon fixation pathways (Hügler and Sievert, 2011) were identified amongst the genomes, with 32 genomes containing ribulose-1,5-bisphosphate carboxylase (RuBisCo) and elements of the Calvin-Benson-Bassham (CBB) cycle, and 7 genomes containing ATP-citrate lyase or citryl-CoA synthetase and citryl-CoA lyase part of the reverse citric acide cycle (rTCA; TABLE 2). A phylogenomic analysis of the 32 putative RuBisCo proteins, identified 5 genomes with type IV RuBisCo and incomplete CBB cycles (SUPPLEMENTARY FIGURE 5).

**Table 2.**
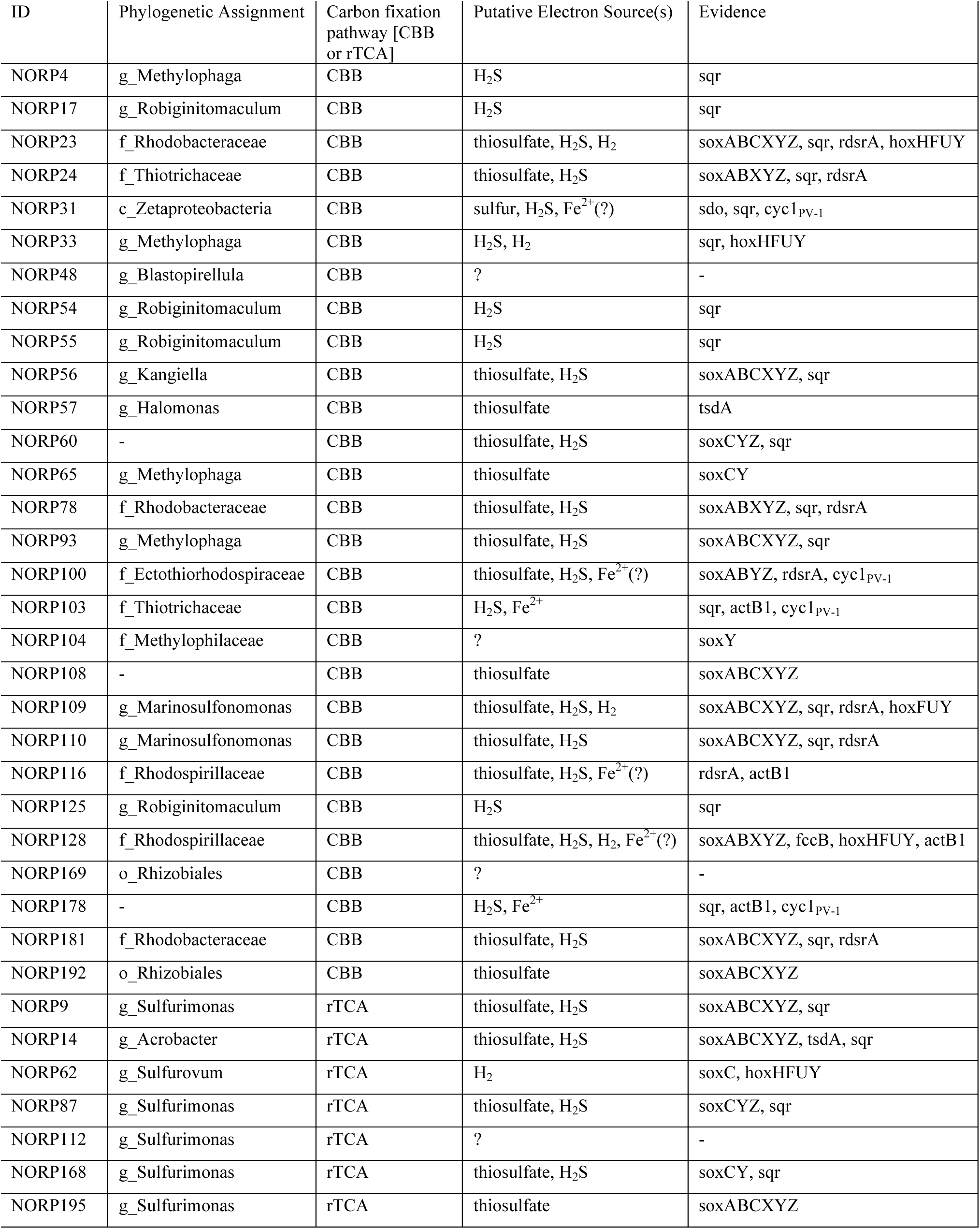
Genomes with carbon fixation potential and possible electron donor sources.

Analysis of the putatively carbon fixing genomes for possible electron donors revealed at least 5 sources: sulfide (HS^-^), sulfur (S^0^), thiosulfate, hydrogen (H_2_), and ferrous iron (Fe^2+^; TABLE 2). The putative electron donors of four of the genomes could not be identified. The most prevalent electron donor as indicated by the presence within the genomes was HS^-^, as 25 genomes had the potential to utilize HS^-^ via either sulfide:quinone reductase (*sqr*) or reverse sulfite reductase (*rdsrA;* SUPPLEMENTARY FIGURE 6). Twenty-one genomes possessed either the components for the SOX system or thiosulfate dehydrogenase (*tsdA*), suggesting a potential for thiosulfate oxidation. The presence of NAD-reducing hydrogenase, capable of the reversible H_2_ redox reactions, offers an avenue for H_2_ as an electron donor in 5 of the genomes capable of carbon fixation. A single genome (NORP31) possessed a sulfur dioxygenase (*sdo*) that could utilize S^0^ as an electron source. Lastly, 6 genomes were identified that may mediate the oxidation of Fe^2+^ linked to carbon fixation, based on the presence of dissimilatory Fe^2+^ molybdopterin oxidoreductase(Tully and Heidelberg, 2016) (Act1B; SUPPLEMENTARY FIGURE 7; SUPPLEMENTARY DATA 6) and/or Fe^2+^ reactive cytochromes (Barco *et al.*, 2015; Cyc1_PV-1_; SUPPLEMENTARY DATA 7).

In assessing the potential for aerobic respiration, 20.5% of the genomes were determined to possess low-oxygen sensitivity (aerobic; aa_3_- and/or bo-type) cytochromes, 12.3% contained high-oxygen sensitivity (microaerobic; cbb_3_- and/or bd-type) cytochromes, and an additional 55.9% contained cytochromes for both aerobic and mircoaerobic oxygen metabolism (SUPPLEMENTARY FIGURE 4; SUPPLEMENTARY DATA 8). An assessment of anaerobic metabolisms showed that 7.7% of genomes possessed the potential to perform complete denitrification (*nirK*/*nirS*, *norBC*, and *nosZ*). An additional 19.5% of genomes were annotated to have the potential to perform a single step in denitrification process (nitrite reduction, nitric oxide reduction, or nitrous-oxide reduction), while 9.2% of genomes could potentially perform only two of the three steps (SUPPLEMENTARY FIGURE 4). Further, one genome (NORP6) contained a ‘true’ sulfite reductase, necessary for complete sulfate reduction (SUPPLEMENTARY FIGURE 6).

### Ecological units and community metabolic function

The genomes that had an appreciable presence (>0.05%, n = 134) in the U1382A samples were placed into 6 ecological units (Unit I-VI) representing 98 genomes that had distinguishing temporal patterns throughout the time series (FIGURE 4; SUPPLEMENTARY FIGURE 8). An additional ecological unit (Unit VII) consisted of eight genomes that were generally cosmopolitan in the U1382A samples. These ecological units represent a progression of community structure over the course of the time series, though in several instances members of an ecological unit re-occur in multiple time points (FIGURE 4). For example, Unit I consists of genomes originally sampled in 2012 that are not observed in TP1, but are observed 12 months later in the TP2 sample. This pattern can also be observed in Unit II (genomes in TP1 seen in TP3-5) and Unit V (genomes in TP2 seen in TP7, 8, and 2014) and supports the results of the sample clusters that indicate that TP1 and TP3-6 are more similar to one another than TP2 and TP7-8 (SUPPLEMENTARY FIGURE 9).

**Figure 4.**
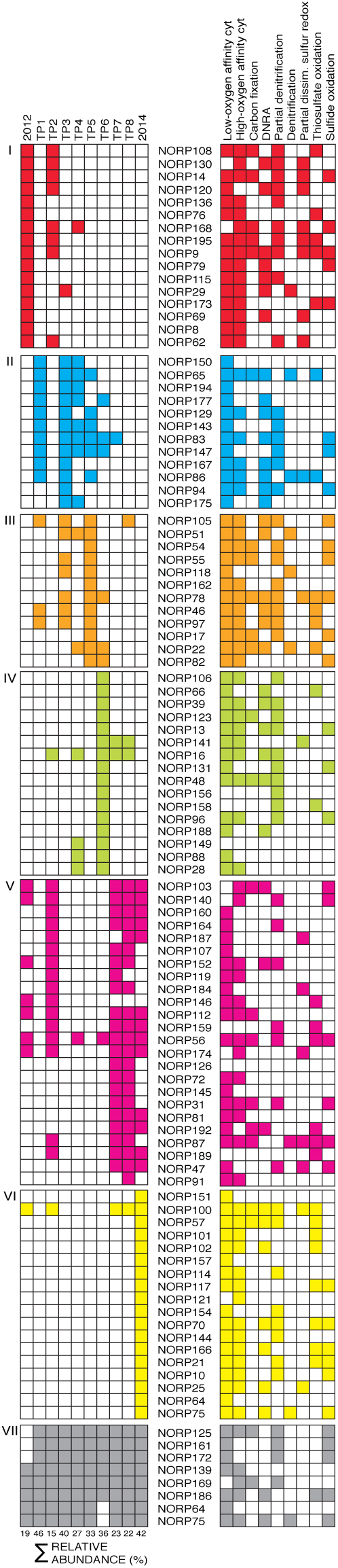
Ecological units and community metabolic function of U1382A. Presence (left column) and predicted function (right column) for genomes assigned to ecological units. Ecological units are ordered to illustrate the progression of community structure through time. Values at the bottom of the column represent the total relative abundance of genomes in ecological units for that time point. Abbreviations: TP, time point; cyt, cytochrome; DNRA, dissimilatory nitrate reduction to ammonia; dissim., dissimilatory

Each ecological unit possessed genomes capable of carbon fixation, partial and complete denitrification, DNRA, thiosulfate oxidation, and sulfur redox, with the exception of complete denitrification in Unit V and sulfur redox in Unit VII. However, analysis of the fraction of the observable community based on relative abundance with a specific functional potential changes considerably over time (FIGURE 5). During the time series, organisms capable of DNRA and complete denitrification are positively correlated (linear regression, R^2^ = 0.86), which is reflected in the fact that most organisms capable of complete denitrification were also capable of DNRA, though the reciprocal was not true (FIGURE 5). This disparity between organisms with DNRA and complete denitrification likely explains why the fraction of DNRA capable community members was always greater than complete denitrification. Similarly, thiosulfate oxidation and sulfur redox processes were positively correlated (linear regression, R^2^ = 0.80), though these functions generally occur in different genomes. The community fraction capable of sulfide oxidation tracks with the observed ecological units and sample clusters, as sulfide oxidation was the most prevalent sulfur pathway in TP1, 3-6, while thiosulfate oxidation was more prevalent in TP2, 7-8, and 2014.

**Figure 5.**
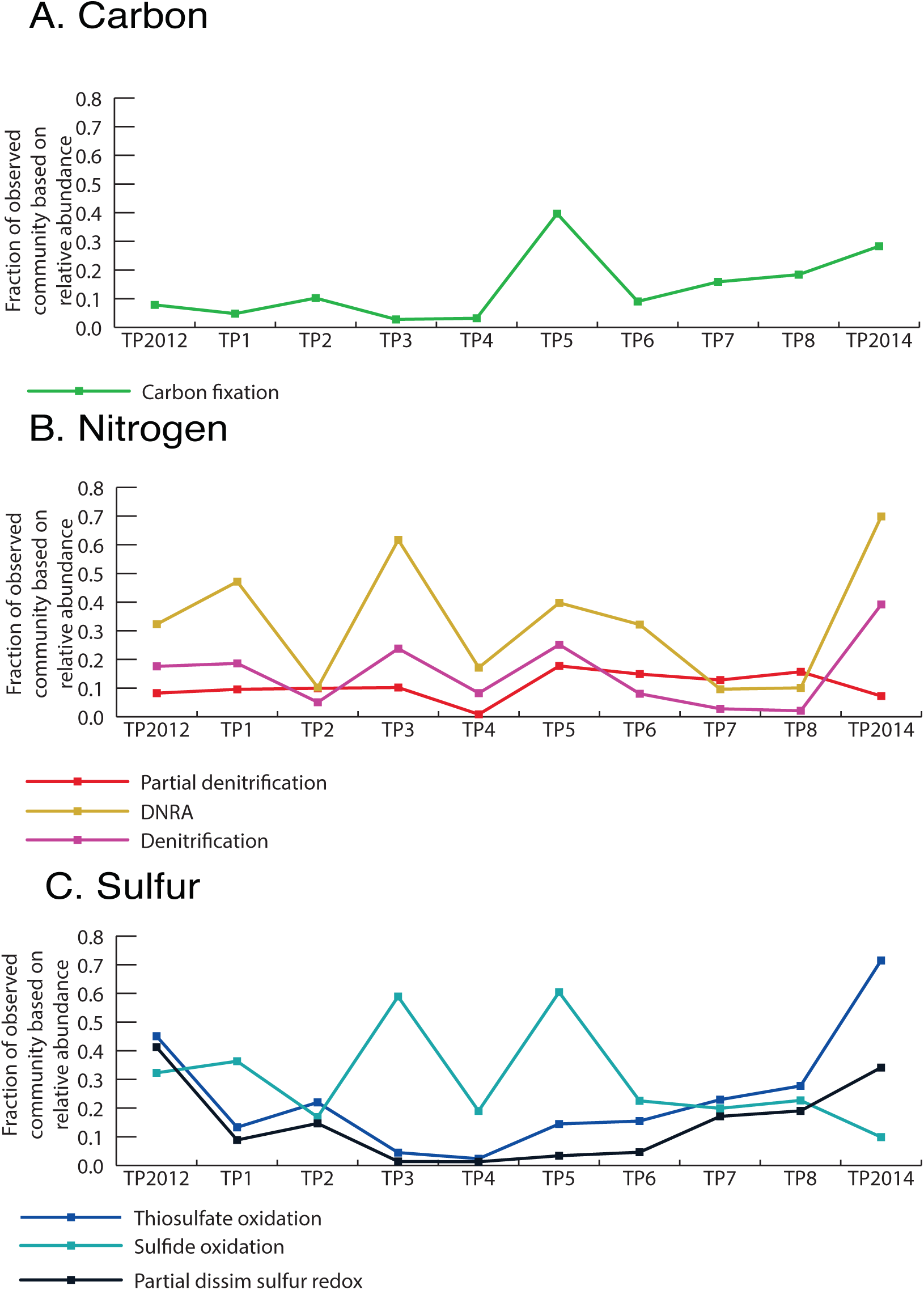
Fraction of the observed microbial community with potential to contribute to biogeochemically relevant processes. **A**. Carbon fixation. **B**. Nitrogen cycing. **C**. Sulfur cycling. Abbreviations: DNRA, dissimilatory nitrate reduction to ammonia; dissim., dissimilatory

Comparison of the potential function of genomes assigned to U1382A, U1383C, and DABW, the number of genomes with a predicted function, and the fraction of the observed community with that function indicates that genomes associated with the DABW do not possess the capability for carbon or nitrogen fixation, denitrification, or sulfide oxidation (FIGURE 6). Based on relative abundance patterns, U1382A and U1383C are dominated by different microbial communities (FIGURE 3), but the number of genomes capable of the various steps in the nitrogen, sulfur, and carbon cycles do not vary (FIGURE 6). Further, there was no statistical difference (Student’s t-test and Wilcoxon rank sum, p < 0.5) between U1382A and U1383C based on the fraction of the observed community capable of an ascribed metabolic reaction, with the exception of ammonia oxidation (Student’s t-test and Wilcoxon rank sum, p = 0.005).

**Figure 6.**
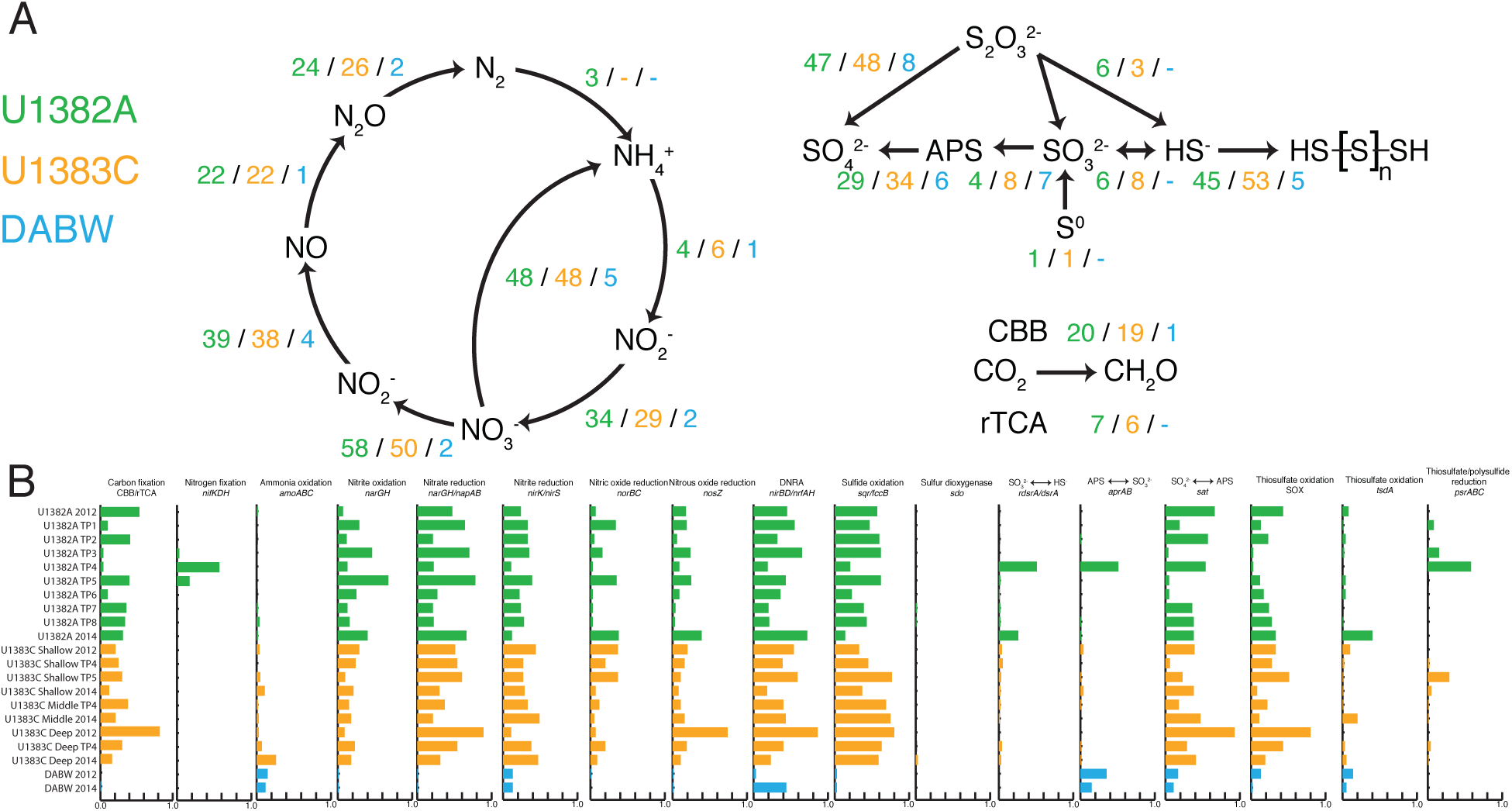
Comparison of putative microbial functionality between U1382A, U1383C, and Deep Atlantic Bottom Water. **A**. Metabolic pathways for nitrogen, sulfur, and carbon fixation, with numbers indicating the number of genomes assigned to a particular sample type with that predicted function. **B**. For each metabolic step represented in A, the fraction of the observed community from each sample that possesses that metabolic step. Abbreviations: TP, time point; DNRA, dissimilatory nitrate reduction to ammonia.

## Discussion

Despite being the largest actively flowing aquifer on Earth, our understanding of microbial communities and their role in biogeochemical cycling in subseafloor crustal fluids is largely unknown. The bulk of our understanding is from studies of fluids from warm environments, including the Juan de Fuca Ridge flank in the NE Pacific Ocean and hydrothermal vents around the globe (Huber *et al.*, 2002; Takai and Horikoshi, 1999; Reveillaud *et al.*, 2016). These environments are characterized by high temperature (25-80°C), low oxygen fluids that are usually dominated by mesophilic and (hyper)thermophilic microorganisms with microaerobic and anaerobic metabolisms (Cowen *et al.*, 2003; Huber *et al.*, 2006; Jungbluth *et al.*, 2013; 2016). This is in contrast to North Pond, which represents a common, but understudied type of ridge flank region, where circulating fluids are cold (4-15°C) and oxygenated (Edwards *et al.*, 2012; Meyer *et al.*, 2016). Previous work at North Pond showed that the fluids in the basaltic crust have similar chemistry to the oceanic bottom water, but that the microbial community has a distinct population structure with potential for both heterotrophic and autotrophic activity (Meyer *et al.*, 2016). Using the increased temporal and spatial sampling offered by our metagenomic time series at North Pond, we verified that the microbial community composition of the crustal fluid samples is fundamentally different from the DABW, and extended this finding to microbial communities and their genomic functional potential using MAGs (FIGURE 2 and 5, SUPPLEMENTARY DATA 10). Further, we also found that the microbial communities within the crustal fluids show shifts in the dominant phyla (and proteobacterial classes) over time within a single hole and between the two holes (FIGURE 2). *Gammaproteobacteria* are dominant in 10 of the crustal fluid samples, but several other phylogenetic groups, *Alpha*-, *Delta*-, *Epsilonproteobacteria*, and *Bacteroidetes*, are abundant in other samples. The initial samples were collected in 2012, approximately six months after the holes were drilled and the CORK systems were installed, therefore it is possible that the observed shifts are due to the holes returning to a natural state after the perturbation of drilling, during which surface water is pumped into the borehole to clear cuttings, inevitably pumping surface waters into the formation. Such shifts in subseafloor crustal fluid community structure have been documented in samples collected shortly after drilling and for several years afterwards on the flanks of the Juan de Fuca Ridge, a younger, warmer crustal system (Jungbluth *et al.*, 2012; 2016), highlighting the importance of time series for understanding such ecosystems and potential stresses. However, the magnitude of chemical shifts observed in discrete samples collected in 2012 and 2014 suggests only minor changes in geochemistry, including a decrease in dissolved oxygen concentrations and increase in dissolved silica concentrations at all four sampling horizons. Increases in dissolved silica may result from either diffusive exchange with sediment pore waters or water-rock reactions at low temperatures, whereas the decrease in oxygen concentrations indicates continued consumption of oxygen (Ziebis *et al.*, 2012; Meyer *et al.*, 2016).

The high resolution analysis, provided by the relative abundance of the reconstructed genomes, reveals that the microbial communities of U1382A, U1383C, and the DABW are composed of distinct MAGs (FIGURE 3). Importantly, genomes from the DABW form a cohesive group of organisms that were not present (or had a limited presence) in the crustal fluids, and conversely none of the crustal-originating genomes were detected in the DABW.

From these results, it is clear that the genomes we reconstructed represent residential subseafloor bacteria and archaea from North Pond crustal fluids, thus allowing for detailed examination of microbial metabolic functions and community dynamics and interactions within the North Pond crustal habitat. It is important to note, however, that the reconstructed genomes only represent a subset of the total microbial community from any one of the metagenomic samples, thus we can only interpret results from the observed community members (SUPPLEMENTARY DATA 4). It is likely, though, that due to the dynamics of assembly and binning that these genomes represent many of the most abundant organisms in the environment. Collectively, the repeated sampling at multiple sites, together with the successful binning of hundreds of genomes, provides an unprecedented dataset for investigation of microbial communities and their associated metabolic potential in the cold, oxic crustal aquifer.

### Carbon fixation

Previous results from North Pond samples in 2012 showed lower concentrations of DOC in the crustal fluids compared to seawater, as well as the potential for carbon fixation, with higher potential rates of autotrophy in the crust compared to seawater, especially at warmer temperatures (25°C) and deeper in the crust (Meyer *et al.*, 2016). In addition, limited metagenomic analysis of three samples from 2012 showed the presence of some genes associated with carbon fixation (Meyer *et al.*, 2016). Our assessment of genomes for the presence of genes representative of autotrophic carbon fixation resulted in the identification of two carbon fixation pathways: the Calvin-Benson-Bassham (CBB) cycle and the reverse citric acid (rTCA) cycle (TABLE 2). All instances of the rTCA cycle were identified within the *Epsilonproteobacteria*, and the CBB cycle was identified in several different groups, including the *Alpha*-, *Gamma*-, and *Zetaproteobacteria*, as well as the *Planctomycetes.* Each of the genomes with potential for carbon fixation was also analyzed for pathways that could provide a lithotrophic source of reducing potential necessary for carbon fixation (TABLE 2). Results indicate that the most prevalent electron source identified amongst the putative carbon fixing genomes was sulfide, but several other electron sources were also identified, including thiosulfate, ferrous iron, sulfur, and hydrogen. These electron sources are likely coupled to the reduction of oxygen, as all but one of the genomes with predicted carbon fixation possess aerobic or microaerobic terminal oxidases. Possible additional terminal electron acceptors include nitrate and the intermediates of denitrification, with all but two of the carbon fixation genomes possess components of the denitrification or DNRA pathways (SUPPLEMENTARY FIGURE 4). While a majority of the genomes with carbon fixation potential are linked to the oxidation of sulfur compounds, a group of genomes have the potential to utilize both H_2_ and Fe^2+^ to drive biomass production in support of the model proposed by Bach (2016). These putative energy couples are congruent with the hypothesis of subseafloor microbial communities that can take advantage of the redox gradient created by the presence of reduced material in volcanic-derived basalt rocks and the oxygenated aquifer fluids(Bach and Edwards, 2003; Bach, 2016). Hydrogen sulfide and iron species have not been detected in the crustal fluids at North Pond (Meyer *et al.*, 2016), but the oxidation of the iron in sulfide complexes in crustal rocks (via biotic or abiotic process) would increase access to sulfide compounds for microorganisms (Barco *et al.*, 2017) and for the abiotic oxidation of sulfide to thiosulfate (Moses *et al.*, 1987). In this manner, it would be possible to sustain carbon fixation through multiple lithotrophic pathways, which are likely important due to the oligotrophic nature of the crustal fluids. This is similar to the prevailing theory in regards to terrestrial crustal systems (Hallbeck and Pedersen, 2008), where lithoautotrophic growth in microorganisms via the CBB cycle has been found in deep terrestrial aquifers in the Fennoscandian shield (Wu *et al.*, 2015).

### Genomic evidence for the prevalence of hypoxic conditions

All measurements at North Pond show that the aquifer fluids at North Pond are oxygenated, with O_2_ concentrations equal to or slightly less (185-244 μM) than that of the DABW (~250 μM; Table 1; (Meyer *et al.*, 2016). Therefore, it was unexpected to find that many of the North Pond genomes had genes that suggest hypoxic or potentially anoxic conditions. More than half of the genomes (56%) had terminal c-type cytochromes for both aerobic (aa_3_- and bo-type) and microaerobic (cbb_3_- and bd-type) metabolisms, with an additional 13% of genomes only possessing the microaerobic cytochromes (SUPPLEMENTARY FIGURE 4). There was substantial evidence that the organisms in this environment were capable of the reduction of nitrate via both dissimilatory nitrate reduction to ammonia (DNRA; 36%) and denitrification (36%; SUPPLEMENTARY FIGURE 4). Further, NORP6 possessed the canonical sulfite reductase, necessary for the anaerobic conversion of sulfite to sulfide (SUPPLEMENTARY FIGURE 4). The role that these genes, commonly associated with anaerobic metabolisms, play in the environment is unclear. It is possible that, similar to sub-oxic microenvironments encountered in the oxic surface ocean (Ploug *et al.*, 1997), the subseafloor hosts microenvironments in which anaerobic metabolisms are ecologically viable. Like the surface ocean, one possible source of such microenvironments may be organic-rich particles, that can be readily colonized by heterotrophic microorganisms. In 2012, samples collected from North Pond crustal fluids showed a high heterogeneity of particles as detected on GFF filters (Meyer *et al.*, 2016). Another possibility may be that the complex and fractured structure of the crustal aquifer provides both oxic and sub-oxic conditions. For example, hydrogeological studies of the Juan de Fuca Ridge flank indicated that fluid flow through the crust likely only occurs through small, discrete channels, restricted to a small volume (<1%) of the crust (Fisher and Becker, 2000). Consequently fluid flow would be highly channelized through a small volume of the crustal rock. While measurements at North Pond CORKs show abundant oxygen, it is possible there are regions where fluid flow slows down and fluids could become stagnant, and anaerobic metabolisms may be more significant to the community as oxygen is consumed by heterotrophic activity or abiotic reactions. However, such stagnant fluids would likely not be indicative of the large crustal flow. Overall, the lack of an appreciable signal in the geochemical data may be the result of the extremely low biomass (~10^4^ cells mL^-1^) and relatively recent entrainment of the formation fluids, especially in U1382A.

### Variable inter- and intra-borehole metabolic diversity

The microbial community observed in U1382A can be effectively assigned to seven ecological units with distinct occurrence patterns (FIGURE 4). These ecological units generally progress in sequential order, though several genomes within an ecological unit were detected in multiple time points, with up to 11 months between samples (TP2 vs. TP7). This re-occurrence of members of the community suggest that there is mechanism for organisms to persist in the aquifer, either locally or transported from elsewhere within the subseafloor. Patterns may also be related to local geochemical conditions, where growth, and thus relative abundance, is tied to specific metabolic processes. Despite these changes in community structure over time, the genomes that are present in the ecological units are functionally redundant, with various metabolisms related to carbon fixation and nitrogen and sulfur cycling present in each of the measured time points (FIGURE 4). While the ecological units as a whole are functionally redundant, the fraction of the observed community capable of a specific metabolic potential shifts over the course of the time series (FIGURE 5A-C). Shifts in genomes capable of nitrate reduction (DNRA and complete denitrification) and sulfur oxidation (thiosulfate oxidation and sulfur redox) processes were positively correlated, suggesting that these metabolic pairs are linked to the same environmental change. Further, shifts in the fraction of the community capable of sulfide oxidation is linked to a microbial community structure that overlaps TP1 and TP3-6, while thiosulfate oxidation is linked to overlaps in TP2 and TP7-8 (FIGURE 5A-C; SUPPLEMENTARY FIGURE 9). This suggests that changes in availability of sulfide and thiosulfate are responsible for the changes in microbial community structure, or conversely, that microbial community metabolic potential impacts the availability of sulfide and thiosulfate.

In comparing U1382A and U1383C, several large, cohesive microbial groups were present in both boreholes (FIGURE 3), with organisms more abundant in U1383C clustering together, to the exclusion of organisms more abundant in U1382A. However, it was common for a group of MAGs to be more abundant in one hole and also have a reduced or minimal abundance in the other hole (FIGURE 3). While this result suggests there is some connectivity between the two subseafloor environments sampled by the CORKs, it is also clear that there are distinct, dominant populations within each hole, likewise there are distinct chemical signatures in both. However, the variation in community structure does not result in differences in metabolic potential, with functional redundancy in all queried processes, except for nitrogen fixation (FIGURE 6). This functional redundancy is further reflected in the fraction of the observed microbial community capable of participating in each metabolic step, with no statistically significant difference between the boreholes, except for ammonia oxidation (FIGURE 6). These results indicate that the observed differences in community structure are not related to carbon fixation or nitrogen and sulfur cycling, and are likely governed by environmental parameters that structure spatially distinct communities with a high degree of functional redundancy. A top-down control on community structure could be susceptibility to viral predation, while a bottom-up control may involve limits in trace nutrients or vitamin availability. Continued analysis of these data and future sampling efforts will help to elucidate the extent of these controls on the microbial community.

## Concluding Remarks

The microbial community in the crustal fluids of North Pond is temporally and spatially dynamic. The putative genomes extracted from our time series reveal a microbial community capable of impacting subseafloor biogeochemical cycles for carbon, nitrogen, sulfur, and iron. These potential functions are redundant as community membership varies in time and space, suggesting that the communities present in both boreholes are poised to utilize the redox potential of the oceanic crust and dissolved organic carbon circulating fluids to drive auto- and heterotrophic growth. Further research will elucidate the extent to which these organisms drive global biogeochemical processes.

## Methods

### Sampling, Cell Quantification, and Chemical Analysis

Crustal fluids were collected from the single horizon at U1382A and from the shallow, middle, and deep horizons in U1383C (Edwards *et al*.) using a mobile pumping system (MPS) designed for microbial sampling from CORK fluid delivery lines as described in Meyer *et al.* (2016) and Cowen *et al.* (2012; FIGURE 1). Deployed with the ROV system, MPS connectors are attached to the CORK wellhead via an umbilical to the hydrological zone of interest within the aquifer. Fluid systems were flushed and allowed to equilibrate before sampling, and dissolved oxygen concentrations were measured during pumping using an Aanderaa sensor (Meyer *et al.*, 2016). In 2012, twelve liters of each fluid sample were filtered on to a 0.22μm Sterivex-GP filter (Millipore) as described in Meyer *et al.* (2016). In 2014, twelve liters of each sample was filtered *in situ* and immediately fixed with RNALater (Ambion), as described previously (Akerman *et al.*, 2013). After sampling in 2012, a battery-powered GeoMICROBE sled was left at each CORK for time-series autonomous sampling of the fluid delivery lines (Cowen *et al.*, 2012). For each filter sample, ~10 liters of fluid were filtered *in situ* and immediately fixed with RNALater. For downstream analysis, ~500 mL of fluid were filtered into two Tedlar bags, one containing 54 mL of 37% formaldehyde for cell enumeration and the other with 4 mL of 10% HCl for inorganic chemistry analyses. Sleds were deployed in April 2012 and recovered in April 2014 with samples collected according to Table 1. Upon sled recovery, filters were transferred to fresh RNALater and stored at −80°C, while all bag samples were stored at 4°C (Cowen *et al.*, 2012). Deep bottom water was sampled in 2012 and 2014 via a CTD at 100 m above the seafloor and filtered in the same manner as the crustal fluids onto Sterivex filters. Total microbial biomass in fluids was enumerated with DAPI (4’,6’-diamidino-2-phenylindole; Sigma) and epifluorescent microscopy (Porter and Feig, 1980). Fluids also were analyzed for dissolved silicon and nitrate using automated colorimetric analysis.

### DNA extraction and sequencing

Total genomic DNA was extracted from the filters using a phenol chloroform method, as previously described (Sogin *et al.*, 2006). DNA was sheared to 175 bp using a Covaris S-series sonicator. Metagenomics libraries were constructed using the Ovation Ultralow Library DR multiplex system (Nugen) following manufacturer’s instructions. Paired-end sequencing was performed on an Illumina HiSeq 1000 at the W. M. Keck sequencing facility at the Marine Biological Laboratory. Raw sequence reads underwent quality control using Cutadapt (Martin, 2011; v.1.7.1; -e 0.08 ‐‐discard-trimmed ‐‐overlap=3) to locate and remove Illumina adapter sequences from both ends of the of the read, followed by quality trimming using Trimmomatic (Bolger *et al.*, 2014; v.0.33; PE SLIDINGWINDOW:10:28 MINLEN:75).

### Ribosomal rRNA identification and relative abundance

From the high-quality paired-end Illumina sequencing reads, 16S rRNA gene fragments were identified using Meta-RNA (Huang *et al.*, 2009; v.H3; -e 1e-10). Putative rRNA fragments and associated mate pairs from each sample were processed through EMIRGE (Miller *et al.*, 2011; 2013; emirge_amplicon.py; −l 113 -i 163 -s 33 -a 32 --phred33) to generate full-length sequences using the SILVA (Quast *et al.*, 2012) SSURef111 reference database (https://github.com/csmiller/EMIRGE). Reconstructed 16S rRNA genes were assigned taxonomy using mothur (v1.34.4) by first aligning the sequences to the SILVA SSURef123 database (align.seqs; flip=T), removing sequences that failed to align, if necessary (remove.seqs), and classifying the sequences (classify.seqs; cutoff=80, iters=1000).

Utilizing the high-quality sequence reads, each set of 16S rRNA sequences was used to recruit reads from the corresponding metagenomic sample, randomly subsampled using seqtk (v1.0-r82; https://github.com/lh3/seqtk) to the size of the smallest library (n = 22,142,100 reads). Reads were recruited using Bowtie2 (v.2.2.5; default parameters) and individual counts of reads per 16S rRNA were determined. Read counts were length normalized and used to calculate the relative abundance of each reconstructed 16S rRNA in the sample (SUPPLEMENTARY DATA 15). Relative abundances were combined for sequences that shared the same mothur-ascribed Phylum (or Class for Proteobacteria) level.

### Metagenomic assembly and binning

High-quality sequence reads were subjected to two rounds of assembly. A primary set of contigs was generated using IDBA-UD (Peng *et al.*, 2012; v.1.1.1; default parameters) utilizing the reads from each individual sample. A secondary set of contigs was generated in Geneious (Kearse *et al.*, 2012) v6.1.8; modified parameters used for “High Sensitivity/Slow”; SUPPLEMENTARY DATA 11) by combining the primary set of contigs ≥500bp in length from samples with the same source (*i.e.*, combining all primary contigs generated from U1382A, etc.). Secondary contigs ≥5kb in length from U1382A and U1383C were combined with secondary contigs ≥3kb generated from DABW (SUPPLEMENTARY DATA 4).

The size-selected set of secondary contigs was used to recruit high-quality sequencing reads from each sample using Bowtie2 (as above). A coverage value, equivalent to recruited reads per bp, was determined for each contig in each sample, the coverage values were log(n + 1) transformed, and subjected to binning using affinity propagation (Frey and Dueck, 2007) and a pre-release version of BinSanity (Graham *et al.*, 2017; -p −1). CheckM (Parks *et al.*, 2015; v1.0.3; lineage_wf) was used to assess the results of the binning utilizing a ≥50% completeness threshold to identify putative genomes. Multiple bins were identified above the completeness threshold that contained substantial estimated contamination (>55% contamination). For each suspect bin, the %G+C and coverage values for each contig were plotted against each other (data not shown) and manually assessed for putative cohesive groups (SUPPLEMENTARY DATA 12).

### Phylogeny

Each putative genome was assessed for the presence of 16 conserved ribosomal marker proteins (Hug *et al.*, 2016) based on a HMMER (Finn *et al.*, 2011) search (v.3.1b2; hmmsearch ‐‐ cut_tc ‐‐notextw) of TIGRfam (Haft *et al.*, 2003) and Pfam (Bateman *et al.*, 2002) models corresponding to the proteins (SUPPLEMENTARY DATA 13). If multiple copies of a ribosomal markers protein were detected, that protein was not included as a marker for that genome, and any genome with <8 markers was not included for further phylogenetic assessment. Ribosomal markers were collected from 1,652 reference genomes representing the major Families and/or Genera from within the Bacteria. Each marker gene from the putative and reference genomes was aligned using MUSCLE (Edgar, 2004; v3.8.31; -maxiters 8), trimmed using trimAL (Capella-Gutiérrez *et al.*, 2009; v. 1.2rev59; -automated1), imported in Geneious (Kearse *et al.*, 2012), and manually assessed and trimmed, if necessary. All of the individual alignments were concatenated and a phylogenetic tree was constructed using FastTree (Price *et al.*, 2010; v2.1.3; -lg -gamma).

Genomes were assessed for the presence of full-length 16S rRNA genes utilizing RNAmmer (Lagesen *et al.*, 2007; v1.2; -S bac -m ssu). The identified rRNA sequences were aligned to the SILVA SSURef123 database using the web-based SINA aligner (Pruesse *et al.*, 2012; default setting, trailing sequences removed from alignment). Aligned rRNA sequences were added to the SSURef123 NR99 ARB tree (Ludwig *et al.*, 2004) using the ARB Parsimony (Quick) tool (default parameters). Members of the ARB tree phylogenetically related to the North Pond genome rRNA sequences were extracted. ARB-based and North Pond genome 16S rRNA sequences were aligned, trimmed, and manually assessed (as above). The final alignment was used to construct a phylogenetic tree with FastTree (-nt -gtr -gamma). Within the genomes, 53 16S rRNAs were identified within 47 genomes. The 16S rRNA gene tree was (SUPPLEMENTARY DATA 14) used to support or refute the assignments provided from the ribosomal marker tree (RMT) and/or CheckM (SUPPLEMENTARY DATA 16).

### Annotation and metabolic analysis

Putative CDS were predicted for the North Pond genomes using Prodigal (Hyatt *et al.*, 2012; v2.6.3; -m -p meta -q) and submitted to the GhostKoala (Kanehisa, Sato and Morishima, 2016) (default parameters; genus_prokaryotes + family_eukaryotes) for annotation using the KEGG (Kanehisa, Sato, Kawashima, *et al.*, 2016) Ontology (KO) system. Based on these KO assignments, genomes were assessed for the degree to which specific pathways and functions were complete in the individual genomes using the information on canonical pathways available as part of the KEGG Pathway Database (updated, Nov 14 2016) and the script KEGG-decoder.py (www.github.com/bjtully/BioData/tree/master/KEGGDecoder).

Beyond specific assignments to pathways and function, for this manuscript several broad functional metabolic categories were identified for genomic bins based on the presence multiple genes. Cytochromes that participate in oxygen chemistry in aerobic organisms were defined as cytochrome c oxidase, aa_3_-type (*coxABCD*) and cytochrome o ubiquinol oxidase (*cyoABCD*), while microaerobic cytochrome metabolism was defined as cytochrome c oxidase, cbb_3_-type (*ccoPQNO*) and cytochrome bd complex (*cydAB*; García-Horsman *et al.*, 1994). Similarly, sulfide oxidation was determined by the presence of sulfide:quinone oxidoreductase (*sqr*), sulfur dioxygenase (*sdo*), and/or sulfite reductase (*dsrA*), when applicable (see below). Additionally, putative thiosulfate oxidation was assessed based on components of the SOX system (*soxABCXYZ*) and/or thiosulfate dehydrogenase (*tsdA*).

Putative CDS annotated as the large subunit of ribulose-1,5-bisphosphate carboxylase (RuBisCo; K01601) were extracted, along with RuBisCo sequences representing previously described major lineages (Tabita *et al.*, 2007; SUPPLEMENTARY DATA 17). The RuBisCo sequences were aligned, automatically trimmed, and used to construct a phylogenomic tree (as above). A similar procedure was applied to identifying molybdopterin oxidoreductases (MOBs), specifically to identify MOBs associated with Fe^2+^ oxidation. Putative MOBs were identified from the Prodigal-derived CDSs via HMMER (hmmsearch, bit score threshold ≥75) using the molybdopterin Pfam (PF00384). Environmental MOBs were aligned with reference sequences (SUPPLEMENTARY DATA 6), automatically trimmed, and used to construct a phylogenomic tree (as above). To differentiate between sulfite reductases present in sulfur reducing organisms and reverse sulfite reductase present in sulfur oxidizing organisms, putative CDS annotated as DsrA (K11180) were used to construct a phylogenomic tree (as above) with reference sequences (Loy *et al.*, 2009) (SUPPLEMENTARY FIGURE 6). An HMM model designed using homologs of Cyc1_PV-1_ identified in neutrophilic iron oxidizing organisms (Tully and Heidelberg, 2016; Barco *et al.*, 2015) was used to search via HMMER (hmmsearch, bit score threshold ≥445) the putative CDSs of North Pond genomes (SUPPLEMENTARY DATA 7).

### Relative abundance and community structure

Contigs composing the North Pond genomes were used to recruit high-quality sequencing reads using Bowtie2 (default parameters) from the subsampled metagenomic samples (as above). Read counts for each genome were determined using featureCounts (Liao *et al.*, 2014; v.1.5.0-p2; -F SAF) and normalized to reads per bp for the full-length of each genome. Length-normalized relative abundance was determined for each sample:

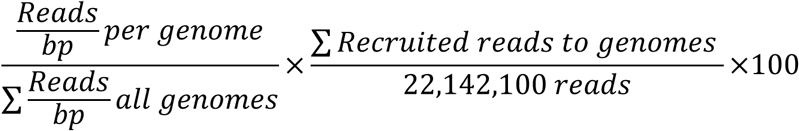

The relative abundance values for the genomes in all 21 sample were used to cluster the genomes and samples in Past3 using the Bray-Curtis similarity measure (SUPPLEMENTARY FIGURE 9). A separate Bray-Curtis clustering step was performed only on genomes detected in the U1382A samples above 0.05% relative abundance (SUPPLEMENTARY FIGURE 8). The observed clusters of genomes were used to determine ecological units that were observed over the course of the time series sampling. Annotations for genomes in the identified U1382A ecological units were used to predict potential function. Genomes from U1382A were assigned to six functions: carbon fixation, partial denitrification (functional assignment predicts incomplete set of denitrification genes), complete denitrification, DNRA, sulfide oxidation, partial dissimilatory sulfur redox (functional assignment predicts incomplete set of sulfur redox genes), and thiosulfate oxidation. The fraction of the observed community with a given function was determined by:

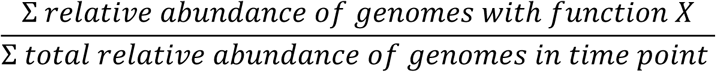

This process was repeated for the complete set of genomes to compare function between U1382A, U1383C, and the DABW. Genomes with relative abundance >0.05% were considered for this analysis and could be assigned to multiple samples. Instead of broad functions, as for U1382A ecological units, genome counts and fractional abundance were assigned based on the presence of key genes in the nitrogen, sulfur, and carbon cycles.

## Acknowledgements

We thank the crews of the R/V Merian and ROV Jason, Wolfgang Bach, Peter Cirguis Chih-Chiang Hsieh, Ulrike Jaekel, Beate Kraft, Huei-Ting Lin, Beth Orcutt, and Heiner Villinger for outstanding support in accomplishing our field programs. Ship time was generously provided by the German Science Foundation (DFG). Both Katrina Edwards and James Cowen’s efforts were critical to the field component and success of this project. They are missed immensely. Leslie Murphy and Emily Reddington provided laboratory support at the W.M. Keck sequencing facility at the Marine Biological Laboratory. This work was supported by NSF OCE1062006 to JAH and NSF OCE1061827 to BTG. The Gordon and Betty Moore Foundation sponsored most of the observatory components at North Pond through grant GBMF1609. The Center for Dark Energy Biosphere Investigations (C-DEBI) (OCE-0939564), a National Science Foundation-funded Science and Technology Centers of Excellence also supported the participation of CGW, JAH, and BJT. This is C-DEBI contribution number XXX.

## Data Availability

[Data submission to NCBI is ongoing. Raw sequences and MAGs will be available under BioProject accession PRJNA391950. Currently, MAGs are available for reviewers via figshare: https://figshare.com/s/939160bb2d41560225581

This project has been deposited at DDBJ/ENA/GenBank under the BioProject accession no. PRJNA391950, drafts of metagenome-assembled genomes are available with accession no. XXX-XXX, and raw sequence reads are available with accession no. XXXX. Raw sequence reads from Meyer *et al*, (2016) constituting the metagenomics samples from 2012, are available under the BioProject accession no. PRJNA280201.

Additional files have been provided and are available through figshare (https://figshare.com/s/939160bb2d4156022558), such as: all primary and secondary contigs; MAGs and bins not analyzed as part of this research; and, all files described as Supplementary Data 1-17.

## Supplemental Information

Supplementary Figures 1-9

Supplementary Data 1-17

Supplementary Figure 1. **Bathymetric map of the North Pond site with CORKs U1382A and U1383C labeled**.

Supplementary Figure 2. **Phylogeny of North Pond putative genomes**. Pseudo-maximum likelihood tree constructed based on the concatenated alignment of 16 conserved ribosomal marker proteins of 140 North Pond and 1,652 reference genomes. North Pond genomes are denoted with stars and an extended terminal branch. Detailed branch lengths and pseudo-bootstrap assignments are provided in SUPPLEMENTARY DATA 5 and inferred phylogenetic assignments are provided in SUPPLEMENTARY DATA 9.

Supplementary Figure 3. **Dendrogram of the Bray-Curtis similarity index from abundance values of 195 genomes in all 21 samples**. Dendrogram structure used orient and cluster genomes in FIGURE 3.

Supplementary Figure 4. **Putative functional assignments for all North Pond genomes**. Heat map displays the fraction of pathway, cycle, and/or gene present in a genome based on KEGG Ontology assignments to biogeochemically relevant processes.

Supplementary Figure 5. **Phylogenetic tree of the large subunit of ribulose 1,5-bisphosphate carboxylase (RuBisCo)**. FastTree generated pseudo-maximum likelihood tree constructed based on an alignment of RuBisCo large subunit from North Pond genomes and previously assigned functional groups.

Supplementary Figure 6. **Phylogenetic tree of subunit A of sulfite reductase**. FastTree generated pseudo-maximum likelihood tree constructed based on an alignment of sulfite reductase subunit A from North Pond genomes and previously assigned functional groups.

Supplementary Figure 7. **Phylogenetic tree of molybdopterin oxidoreductases with various assigned functions**. FastTree generated pseudo-maximum likelihood tree constructed based on an alignment of molybdopterin oxidoreductases from North Pond genomes and previously assigned functional groups.

Supplementary Figure 8. **The relative abundance of genomes identified in the U1382A samples, clustered on dendrogram based on the Bray-Curtis similarity index**. The U1382A samples are ordered based on the relative positions as presented in Supplementary Figure 9. Branches are colored based on which ecological unit a genome was assigned to in Figure 4.

Supplementary Figure 9. **Dendrogram of the similarity between the metagenome samples**. Samples are clustered using the Bray-Curtis similarity index based on the genome relative abundance profile, as presented in Figure 3. Color scheme is to highlight the 2 clusters of samples from U1382A.

Supplementary Data 1. Assembly statistics for the individual samples, as assembled by IDBA-UD, and the combined assemblies, assembled by Geneious.

Supplementary Data 2. FASTA file of the EMIRGE assembled full-length 16S rRNA genes.

Supplementary Data 3. Statistics of the 195 analyzed genomes, including estimated percent completeness and contamination/redundancy.

Supplementary Data 4. The percentage of reads that recruit back to a subset of contigs.

Supplementary Data 5. Newick file containing the 16 ribosomal marker tree used to construct SUPPLEMENTARY FIGURE 2.

Supplementary Data 6. Functional assignment for each molybdopterin oxidoreductase, reference and environmental used to construct SUPPLEMENTARY FIGURE 7.

Supplementary Data 7. Hidden Markov Model used to search for cytochrome *c*_PV-1_.

Supplementary Data 8. The fractional output for each metabolic process searched by KEGG-decoder.py.

Supplementary Data 9. Percent relative abundance determined for each genome in each sample. Used to construct FIGURE 3.

Supplementary Data 10. Relative abundance and taxonomic assignment for full-length 16S rRNAs as reconstructed by EMIRGE.

Supplementary Data 11. Parameters used in the Geneious assembly.

Supplementary Data 12. Methodology and results for splitting the high contamination bins, prior to analysis.

Supplementary Data 13. Hidden Markov Models used to search for the 16 ribosomal markers used in SUPPLEMENTARY FIGURE 2.

Supplementary Data 14. Newick file containing the 16S rRNA tree based on sequences extracted from the assembled genomes.

Supplementary Data 15. Cumulative relative abundance values for phylogenetic groups based on EMIRGE reconstructed full-length 16S rRNA and mothur taxonomy, as determined for FIGURE 2.

Supplementary Data 16. Phylogenetic assignment for each genome, listing results from CheckM, the ribosomal marker tree, and 16S rRNA tree. Conflicting assignments are noted.

Supplementary Data 17. Assignments and accession numbers for the ribulose-1,6-bisphosphate carboxylase reference sequences used to construct SUPPLEMENTARY FIGURE 5.

